# Synthetic genetic circuits enable reprogramming of plant roots

**DOI:** 10.1101/2022.02.02.478917

**Authors:** Jennifer A. N. Brophy, Katie J. Magallon, Kiril Kniazev, José R. Dinneny

## Abstract

The shape of a plant’s root system influences its ability to reach essential nutrients in the soil and to acquire water during drought. Progress in engineering plant roots to optimize water and nutrient acquisition has been limited by our capacity to design and build genetic programs that alter root growth in a predictable manner. Here, we construct synthetic genetic circuits to control gene expression with high spatial precision across root tissues. These circuits produce specific patterns of gene expression by combining the activity of multiple native promoters using logical operations. We then use the circuits to predictably alter root structure. This work demonstrates the ability of synthetic genetic circuits to control gene expression across tissues and offers an exciting means to reprogram plant growth.

## Main Text

Recent advances in our understanding of the molecular mechanisms driving plant development have identified key regulators of important agronomic traits, but our limited ability to control gene expression in plants is a barrier to applying this knowledge (*1*–*4*). Indeed, it remains challenging to express genes in specific patterns in plants, especially if those patterns are outside the reach of native promoters (*5*–*9*). One attractive solution is synthetic genetic circuits, which have been implemented in a wide array of prokaryotic and eukaryotic cell lines (*10*–*14*). Circuits provide a powerful means of controlling gene expression, but circuit technology has been difficult to implement in plants because of the long timescales required for producing transgenic lines and the difficulty of tuning circuit activity across heterogenous cell types (*15*). Here, we show that robust circuits can be built in plants when a quantitative transient expression assay is used to tune circuit components. These circuits can be transferred to whole plants in order to control spatial patterns of gene expression across root tissues and to predictably modify plant development.

To construct logic gates in plants, we first generated a collection of synthetic transcriptional regulators that can be used to control gene expression. We designed simple transcriptional activators, composed of bacterial DNA binding proteins, VP16-activation domains, and SV40 nuclear localization signals (NLS), and synthetic repressors, composed of only DNA binding proteins and NLSs (Fig. 1A, B) (*16*–*19*). Activatable promoters were created by fusing multiple copies of the DNA sequence (operator) that these transcription factors (TFs) bind to, and a minimal plant promoter (−66 to +18 of the Cauliflower Mosaic Virus (CaMV) 35S promoter) (Fig. 1A) (*20*, *21*), while repressible promoters were built by placing one operator sequence at the 3’ end of a full length CaMV 35S promoter (Fig. 1B) (*22*).

**Fig. 1.**
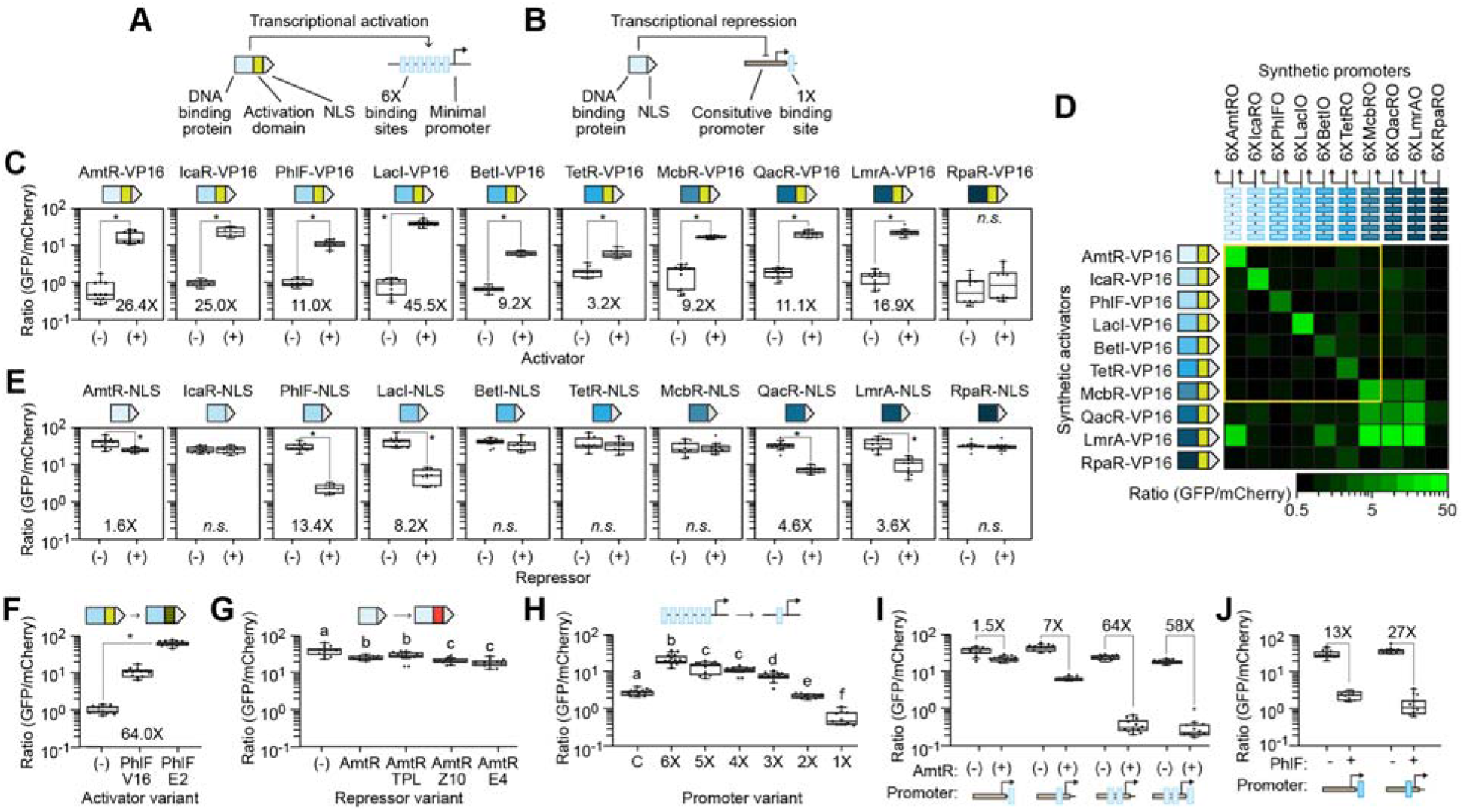
Basic building blocks for constructing synthetic genetic circuits. (**A-B**) Schematics of the synthetic transcriptional activators (A) and repressors (B) built to control gene expression in plants. Activity (**C**) and specificity (**D**) of the synthetic activators. Yellow box highlights orthogonal activators. (**E**) Activity of the synthetic repressors. (**F**) Comparison of PhlF-based activators containing either the VP16 (V16) or ERF2 (E2) activation domain. (**G**) Comparison of AmtR-based repressors containing either no repressive domain, or the TOPLESS (TPL), ZAT10 (Z10), or ERF4 (E4) repressive domain. (**H**) Activity of synthetic activatable promoters containing one (1X) to six (6X) AmtR operators co-infiltrated with AmtR-VP16. Control construct (C) is *proGO190::GFP*. (**I-J**) Engineered synthetic promoters’ response to AmtR-NLS (I) and PhlF-NLS (J) repressors. In all panels, expression is measured in *Nicotiana benthamiana* leaves. TFs are expressed using the constitutive G1090 promoter. Box plots show the median of twelve leaf punches collected from three leaves that were infiltrated and measured on different days. Box plot hinges indicate the first and third quartiles. Dots show individual data points. Stars denote significant differences in normalized GFP expression and letters denote significance groups (p Ã 0.01, student’s two-tailed T-test). *N.s*. not significant. Fold change for significantly active transcription factors was calculated by dividing the average ON state by the average OFF state.

Synthetic TF activity was measured in plants by delivering DNA encoding the proteins and their target promoters to *Nicotiana benthamiana* leaves using *Agrobacterium tumefaciens* (Supplementary Fig. S1). Nine of the ten synthetic activators were able to turn on gene expression at their target promoters in plant leaves (3 to 45-fold increase in expression relative to promoter-only experiments) and seven were specific to the promoters they were designed to regulate (Fig. 1c, d). The specificity of these synthetic activators is important for constructing logic gates free of cross talk between components (*23*). In contrast to the activators, only four of the synthetic repressors generated more than a two-fold change in gene expression (Fig. 1E). Thus, many of the repressors required further optimization before they could be used to control expression in plants.

To achieve greater control over expression, we modified the regulatory domains of the synthetic TFs. Replacing the human herpes virus-derived VP16 activation domain with that of the Arabidopsis ETHYLENE RESPONSE FACTOR 2 (ERF2_AD_) improved activity of the weakly active PhlF-based synthetic activator from 11- to 64-fold (Fig. 1F) (*24*). In contrast, adding repressor domains to the AmtR-based synthetic repressors either had no effect (TOPLESS (TPL^RD^)) or only modestly increased repression from 1.5-fold (NLS only) to 1.8-fold (ETHYLENE RESPONSE FACTOR 4 (ERF4^RD^)) or 2-fold (SALT TOLERANCE ZINC FINGER (ZAT10^RD^)) (Fig. 1G) (*25*, *26*); thus, swapping the activation domain led to increased expression, while adding repressive domains had a more modest effect.

We also engineered the synthetic promoters to tune expression levels and improve dynamic range. Changing the number of TF binding sites in an activatable promoter resulted in a collection of promoters of varying strength. Synthetic promoters with three or more operators amplified expression relative to the constitutive promoter used to drive expression of the activator (compare 6X – 3X with control, Fig. 1H). In contrast, synthetic promoters with two operators roughly replicated the constitutive promoter’s expression level and one operator produced less expression (Fig. 1H). These data suggest that synthetic promoters can be used to tune gene expression levels by either amplifying or dampening the transcriptional signal produced by an input promoter.

In line with a previous study, we found that moving the location of the operator within a repressible promoter improved repression and dynamic range (Fig. 1I) (*14*). Adding a second operator further improved repression and increased dynamic range to 64-fold (Fig. 1I). However, adding a third binding site reduced the promoter’s ON state without significantly reducing the repressed state, resulting in a smaller dynamic range than the two-operator promoter (58-fold) (Fig. 1I). These design features were transferrable to other synthetic repressor-promoter pairs (Fig. 1I, Supplementary Fig. S2). Thus, the number and location of operators are useful design levers that impact promoter activity.

Using these synthetic regulators, we constructed simple circuits that perform Boolean logic operations. Synthetic TFs built with the AmtR and PhlF DNA binding proteins served as the inputs to all circuits and GFP served as the output. Circuit activity was measured in *N. benthamiana* leaves, which were infiltrated with multiple Agrobacterium strains, each containing one plasmid that encoded either an input TF or the output (Fig 2A, B).

**Fig. 2.**
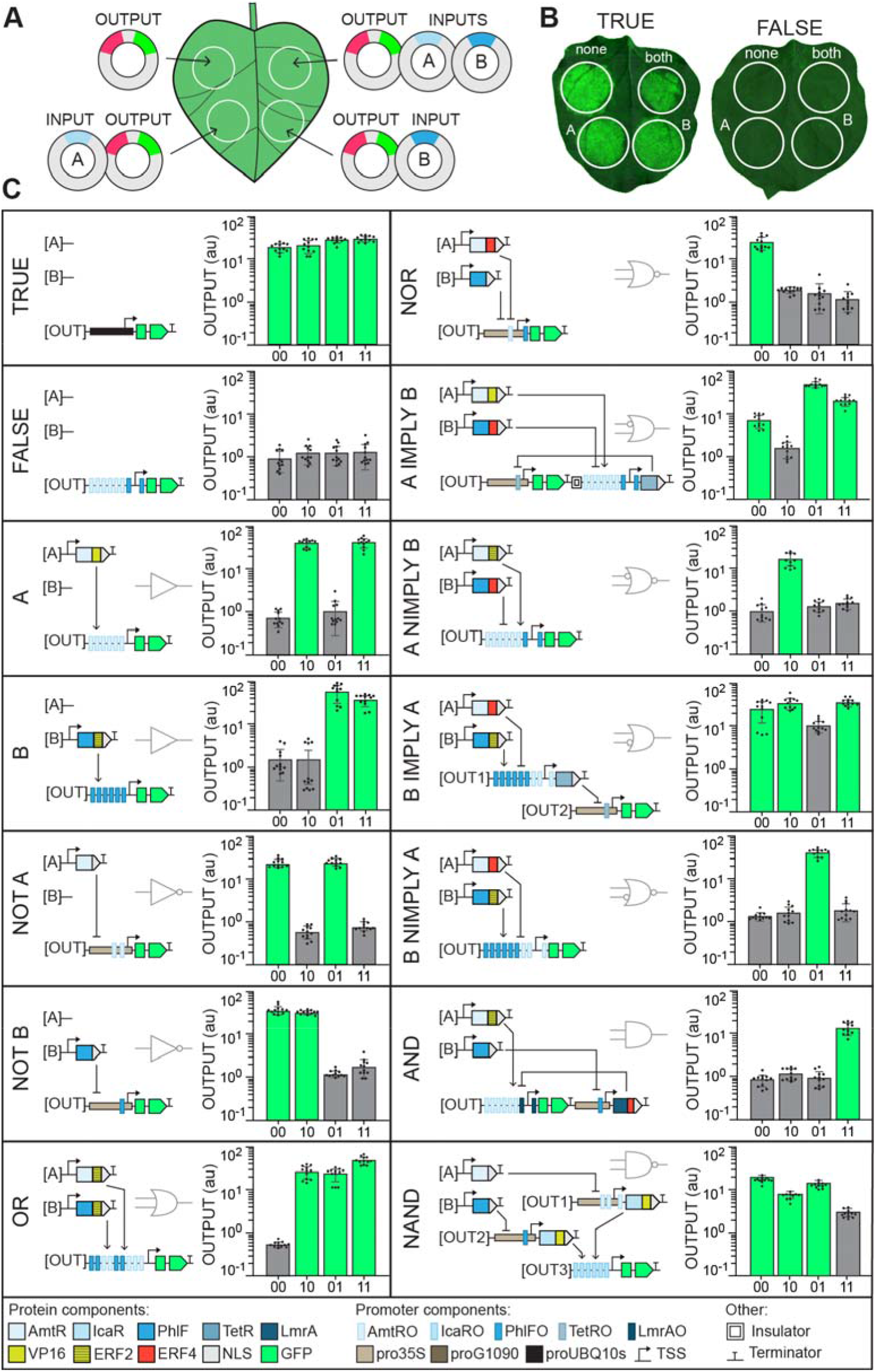
Logic gates in *Nicotiana benthamiana* leaves. (**A**) Schematic for testing circuit activity in *Nicotiana benthamiana* using a transient expression assay (Methods). *N. benthamiana* leaves were infiltrated with Agrobacterium strains containing plasmids that encode either one input transcription factor (A or B) or the output gene *GFP*. mCherry was encoded on the output plasmid and used to normalize fluorescence measurements. (**B**) Example images of leaves infiltrated with TRUE and FALSE gate components. Circles encompass areas of infiltration. (**C**) Gate behavior in *N. benthamiana*. Bar charts show output of each circuit, reported as the ratio of GFP to mCherry fluorescence. Green bars indicate gate states that should be ON and grey bars indicate states that should be OFF to implement correct logic. Data are mean and s.d. of twelve leaf punches collected from three leaves infiltrated and measured on different days. Dots show individual data points. Plasmid sequences and data for individual GFP and mCherry channels provided as Supplementary Data.

In each circuit, computation was performed by synthetic promoters that respond to the TF inputs in unique ways. Simple gates, like the A and B BUFFER gates, use synthetic promoters that are activated or repressed by only one of the input TFs, while more complex gates required synthetic promoters that respond to multiple inputs (Fig. 2C, Supplementary Fig. S3). For example, the OR gate output promoter contains AmtR and PhlF operators, such that it is activated by either TF (Supplementary Fig. S3). For composite promoters to function correctly, both the promoter architecture and activation/repression domains on the synthetic TFs needed to be optimized. The number and arrangement of AmtR and PhlF operators in the OR output promoter affected fold change and relative ON states, with the best design containing alternating pairs of operators (Fig. 2C, Supplementary Fig. S3). For the NOT IMPLIES (NIMPLY) gates, input repressors needed repressive domains, such as ERF4^RD^, to stop synthetic activators from initiating transcription at the output promoter and implement NIMPLY logic (Fig. 2C, Supplementary Fig. S3). However, this requirement could be overcome by increasing the number and location of repressor operators within the synthetic promoter (Supplementary Fig. S3). Most circuits involved several design-build-test cycles, which were facilitated by the rapid *N. benthamiana-based* assays and the modular nature of the synthetic biology parts generated here.

Layered logic gates in which AmtR and PhlF do not directly control expression of GFP, but instead modulate the expression of other synthetic TFs, occasionally encountered problems where expression cassettes were insufficiently insulated from each other. The A IMPLY B gate, which worked well when its output genes were encoded on separate plasmids, had an erroneously reduced ‘no input’ state when both output genes were encoded on the same plasmid (Fig. 2C, Supplementary Fig. S3). In an attempt to fix this problem, we added one or two copies of the previously characterized insulator *Arabidopsis thaliana* MATRIX ATTACHMENT REGION 10 (MAR10) between the two A IMPLY B output genes (*27*). However, insulation only improved gate activity two-fold (Supplementary Fig. S3, Version 2 *vs*. Versions 3 & 4). Consequently, the other IMPLY gate (B IMPLY A) was built with two output plasmids that were co-delivered to plant cells (Fig. 2C).

Functional plant circuits were transferred to model plant *Arabidopsis thaliana* to test their capacity to generate new spatial patterns of gene expression across tissue layers in the root tip. The tissue specific promoters of *SOMBRERO* (*proSMB*, expressed in the entire root cap) and *PIN4* (*proPIN4*, expressed in columella root cap and stele) were used to drive expression of our input TFs (Fig. 3A, B) (*28*, *29*). By combining the activity of these input promoters using logic gates, we expected several different spatial patterns of gene expression (cartoons, Fig. 3C).

**Fig. 3.**
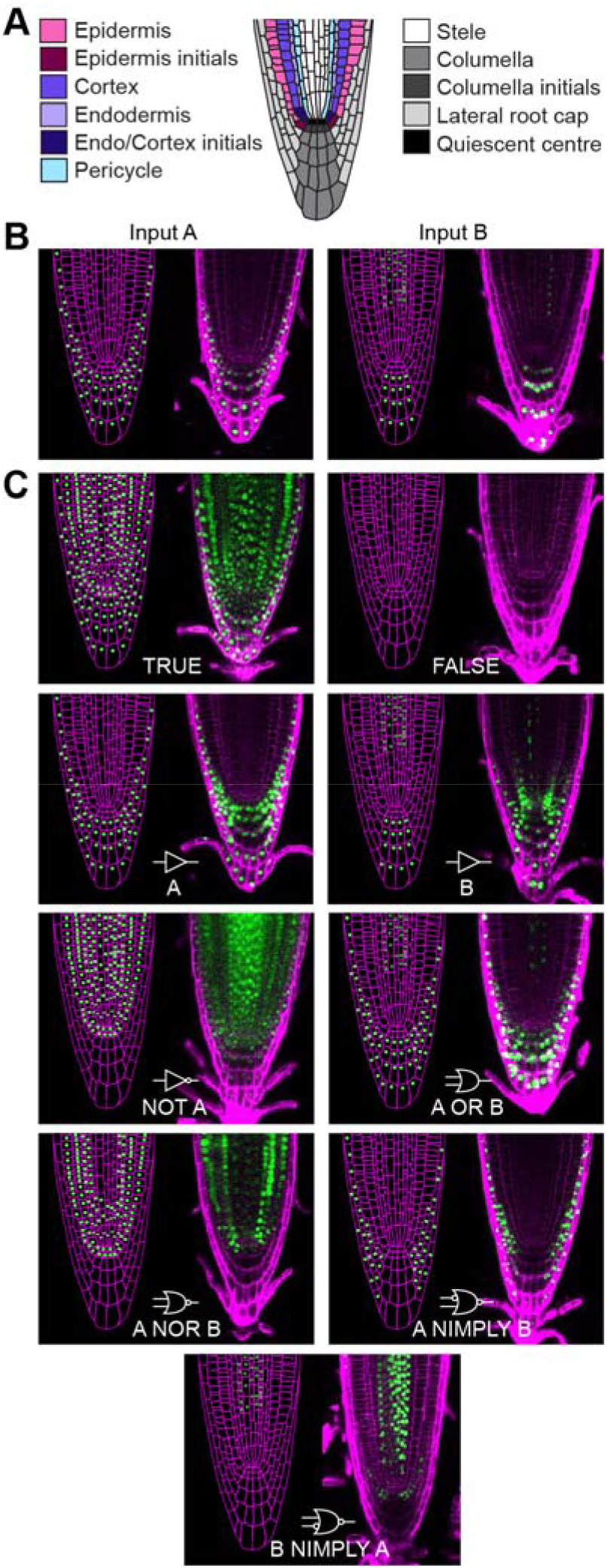
Patterning gene expression using logic gates. (**A**) Cell types in the Arabidopsis root tip. (**B**) Schematics (left) and confocal images (right) showing the expression pattern of input promoters *SOMBRERO (SMB*) (A) and *PIN-FORMED 4 (PIN4*) (B). (**C**) Expected expression patterns (left) and confocal images (right) of logic gates that use the *SMB* and *PIN4* promoters to drive expression of input TFs. Output of each circuit is nuclear localized GUS-GFP fusion protein. All root images were taken five days after sowing. Additional independent lines and T-DNA schematics in Supplementary Fig S4.

We tested nine different logic gates in Arabidopsis, seven of which generated the expected expression pattern (Fig. 3C, Supplementary Fig. S4). For example, the A BUFFER gate faithfully replicated input promoter A’s expression pattern (*proSMB*), while the OR and NOR gates combined and inverted the input promoters’ expression patterns correctly (Fig. 3C, Supplementary Fig. S4).

One notable difference in circuit performance between Arabidopsis and *N. benthamiana* was revealed by the A NIMPLY B gate. In *N. benthamiana*, synthetic repressors containing the ERF4 repressive domain created a tighter OFF state than the simpler PhlF-NLS repressors. In Arabidopsis, the A NIMPLY B gate pattern was incorrect when the gate was built using an ERF4^RD^-containing repressor. With PhlF-ERF4^RD^, the A NIMPLY B gate, which is supposed to be ON in the entire root cap except the columella, was only ON in the outermost root cap layers (Supplementary Fig. S4). This may be due to the activity of the repressive domain, which is believed to recruit histone modifying enzymes that maintain a repressed state in cells that are not actively expressing the repressor (*26*). Thus, in instances where repressors containing the ERF4^RD^ domain generated a logical fault, successful gates were built with PhlF-NLS repressors that lack the capacity to modify histones (Fig. 3C, Supplementary Fig. S4).

The two failed gates (B BUFFER and B NIMPLY A) were both the result of an unexpected pattern created by expressing the AmtR-activator using the Input B (*PIN4*) promoter (Fig. 3C, Supplementary Fig. S4). The B BUFFER gate was expected to directly replicate the PIN4 promoter input, but instead it produced a spatially expanded pattern with expression in the quiescent center (QC) and neighboring initial cells (Fig. 3C, Supplementary Fig. S4). Although the B gate activity does not match our PIN4 promoter control (Fig. 3B, *Input B*), previous studies have found the PIN4 promoter to be active in these cell types (*30*, *31*). Like the B BUFFER gate, B NIMPLY A is aberrantly ON in the quiescent center (QC) and neighboring initial cell types (Fig. 3C, Supplementary Fig. S4). Since B NIMPLY A relies on the same PIN4 promoter and AmtR-activator to turn on the output promoter, the same mechanism that caused a mismatch between the expected and observed B gate pattern may be responsible for this gate’s error. Although the QC and columella/epidermis/endodermis initial expression was unexpected, the B NIMPLY A gate’s logic functioned correctly in Arabidopsis, which is evident by the absence of GFP expression in the lateral root cap. Though well-tuned and functional in *N. benthamiana*, B and B NIMPLY A gates’ aberrant activity in Arabidopsis roots highlight the challenge of implementing circuits that must function in all cell types in heterogeneous tissue.

To demonstrate how precise spatial control over gene expression may be used to engineer development, we modified a key aspect of Arabidopsis root structure: lateral root branch density. Lateral roots enable plants to radially sample soil and the number of lateral roots that a plant generates affects its ability to search for water and essential nutrients in the environment (*32*).The close relationship between root structure and plant fitness has led to proposals of ideal root structures for plant growth in specific environments (called root ideotypes) (*33*). Ideotype hypotheses have been difficult to test directly because genetic changes that affect root structure often have pleiotropic effects. For example, a gain of function mutation in the developmental regulator *INDOLE-3-ACETIC ACID INDUCIBLE 14* (*IAA14*) called *solitary root* (*slr-1*)eliminates root branching, but also hinders root gravitropism, root hair development, and primary root growth (Fig. 4, Supplementary Fig. S5, *slr-1*) (*34*). To disentangle root branching from other developmental processes, we expressed the *slr-1* mutant gene using a tissue specific promoter that is only ON in lateral root stem cells, *proGATA23*^23^. When expressed from *proGATA23*, *slr-’s* impact on development is restricted to lateral roots. In these plants, no lateral roots form, but gravitropism, root hair development, and primary root growth are normal (Fig. 4, Supplementary Fig. S5, *proGATA23::slr-1*).

**Fig. 4.**
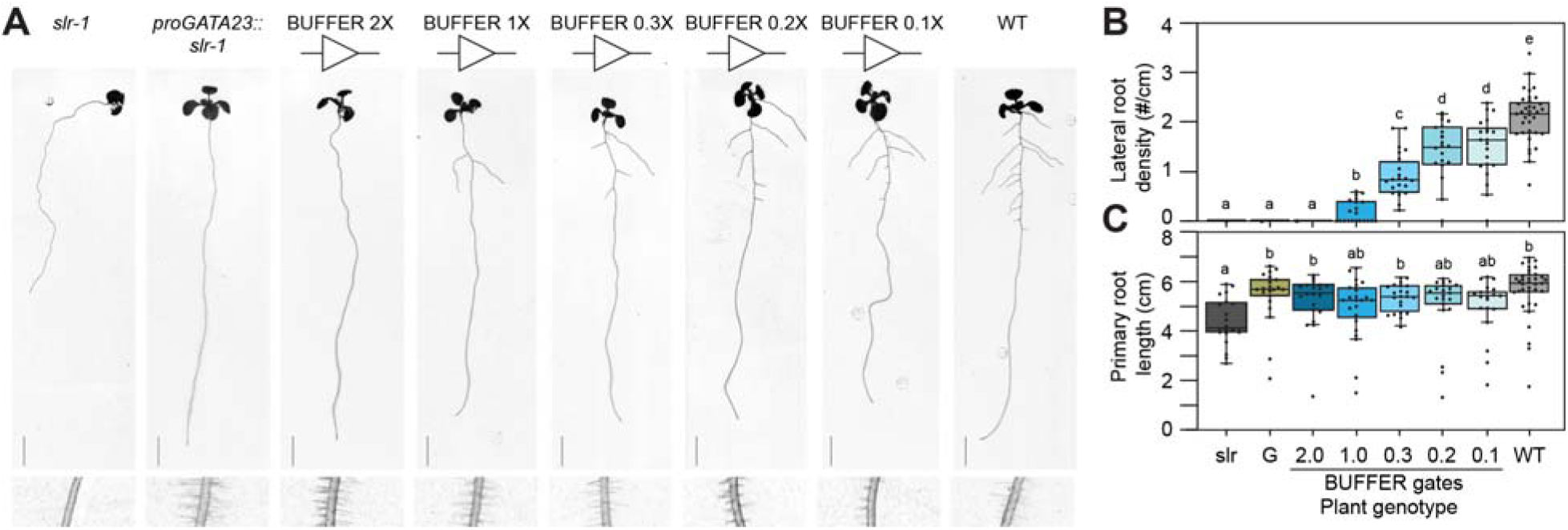
Engineering root branch density using BUFFER gates. (**A**) Root structure (top) and root hairs (bottom) of Arabidopsis plants engineered to modify root branch density. BUFFER gate output promoters contain either one (1X) or two (2X) copies of the consensus AmtR operator or one copy of a mutated AmtR operator (0.1 – 0.3X). Scale bar, one cm. (**B**) Lateral root density calculated as the number of emerged lateral roots divided by total primary root length. (**C**) Primary root length measured from root-hypocotyl junction to the root tip. Control samples are labeled as follows: *slr-1* (slr), *proGATA23::slr* (G), wild type (WT). In (B) and (C), box plots show the median of at least 20 T1 plants. Box plot hinges indicate the first and third quartiles. Letters denote significant differences in LR density or primary root length (p Ã 0.01, Student’s two-tailed t-test). All measurements and images were taken ten days after sowing.

We designed BUFFER gates to express *slr-1* at varying levels in lateral root stem cells to determine whether lateral root branch development could be quantitatively controlled. These BUFFER gates use the *GATA23* promoter to drive expression of the AmtR-VP16 synthetic TF, which then activates a synthetic promoter with one, two, four, and six copies of the AmtR operator to drive expression of *slr-1*. Since plant transformation randomly inserts transgenes into the genome, a single construct can create a range of expression. Although this variation potentially alleviates the need for promoters of varying strength, none of the plants (> 20 independent lines per construct) containing *proGATA23::slr-1* or a BUFFER gate with two, four, and six copies of the AmtR operator produced lateral roots, suggesting that the range of expression conferred by these constructs was above the threshold for fully blocking lateral root development (Fig. 4A, B, data not shown for 4X and 6X promoters). To further reduce the strength of the output promoter, we mutated key residues in the AmtR binding site (Supplementary Fig. S6) (*35*). Weaker synthetic promoters resulted in BUFFER gates that produced a gradient of lateral root densities, roughly correlated with the strength of the synthetic promoter (0.1 - 1X, Fig. 4). Importantly, none of the BUFFER gates affected primary root growth, root hair development, or gravitropism (Fig. 4B, C, Supplementary Fig. S5). Thus, BUFFER gates enabled tissue specific tuning of expression level and the results demonstrate the importance of controlling both expression level and patterns for making targeted changes in root structure.

Our work demonstrates that synthetic genetic circuits can be used to program gene expression across cell types in plants. This approach expands the impact that a handful of characterized promoters can have on our ability to express genes in specific cells and presents an alternative to the time-consuming process of searching genomes for tissue specific promoters with desired expression patterns. The ability to quantitatively control gene expression levels in a tissue specific manner presents an opportunity to probe gene dosage effects in a tissue specific manner, which previously lacked a reliable framework in plants. Additionally, the framework for circuit construction we have developed could be applied to reprogram plant responses to the environment in a tissue specific manner. As climate and agricultural challenges become more formidable, genetic programming capabilities enabled by synthetic genetic circuits should become increasingly utilized for the introduction of novel traits into plants.

## Materials and Methods

### Bacterial strains and growth conditions

*E. coli* strain NEB10β (*Δ(ara-leu) 7697 araD139 fhuA ΔlacX74 galK16 galE15 e14-⍰80dlacZΔM15 recA1 relA1 endA1 nupG rpsL (StrR) rph spoT1 Δ(mrr-hsdRMS-mcrBC)*) was used to construct all plasmids. *Agrobacterium tumefaciens* strain GV3101 containing helper plasmid pSoup (*TetR*) was used for all plant transformations (36).

All bacteria were grown in LB medium (Sigma L3022) in incubating shakers at 250□r.p.m. (New Brunswick Scientific Innova 42). *E. coli* strains were grown at 37□°C and *A. tumefaciens* strains were grown at 28□°C. Antibiotics were used at the following concentrations: carbenicillin 100□μg/ml (Gold Bio C-103), kanamycin 50□μg/ml (Sigma K4378), spectinomycin 100□μg/ml (Sigma S4014), gentamycin 50□μg/ml (Corning 30-005), rifampicin (Gold Bio R-120), and tetracycline 10□μg/ml (Sigma T7660).

### Plant growth conditions

*N. benthamiana* plants were grown and maintained in a Percival growth chamber at 25°C in 12/12-h light/dark cycles with 60% humidity. Plants were grown in PRO-MIX PGX soil (Pro Mix 30380RG). Light was provided by four far red bulbs and white fluorescent lights at a total intensity of ~100 microEinsteins.

Arabidopsis plants were grown and maintained in Percival-Scientific growth chambers (CU41L4) at 22°C in 16/6-h light/dark cycles with 60% humidity. Light was provided by LED arrays at a total intensity of ~100 microEinsteins. Plants for transformation were grown in 4-inch pots and PRO-MIX PGX soil (Pro Mix 30380RG). Plants for imaging and root structure analysis were plated as seeds onto sterile 0.7% Gelzan (Sigma G19010) media containing 1X Murashige and Skoog nutrients (Caisson MSP01-50LT) and 1% sucrose (Sigma S0389) at pH 5.7. Seeds were surface sterilized prior to plating with 95% ethanol and 20% bleach. Plates were sealed with micropore tape (3M 1530-0) to allow for gas exchange.

### Transgenic plant lines

Transgenic plants were generated by Agrobacterium tumefaciens-mediated floral dip of Arabidopsis thaliana Columbia ecotype (37). Briefly, *A. tumefaciens* cultures were inoculated from frozen into 14-ml culture tubes (Fisher Scientific 352059) containing 2 mL of LB media with antibiotics. Cultures were grown for 24 h at 28°C and 250 r.p.m. After incubation, 250 μL of bacterial culture was plated on 15 cm round LB agar plates containing antibiotics and incubated for two nights at 28°C. Next, the cells were scraped from plates and resuspended in 200 mL transformation buffer (per Liter: 50 g sucrose (Sigma S0389), 250 μL silwet-77 (Lehle Seeds VIS-30), 0.93 g MgCl_2_ (Sigma M4880)). Arabidopsis inflorescences were submerged in the bacterial solution for 2 minutes and then transferred to a dark, humid container where they were allowed to incubate overnight. The following day, transformed plants were returned to normal growth conditions where they continued to grow until they produced seeds.

Transgenic seeds were isolated using fluorescence microscopy. All of the plasmids in this study contain a constitutively expressed mCherry marker (*proUBQ10::mCherry*) (similar to reference (38)). This marker enables selection of T1 transgenic plants by screening for red fluorescence in dry seeds.

### Plasmid Construction

All constructs in this study were generated using a mixture of Gibson and Golden Gate cloning methods (39, 40). Most DNA parts (promoters, CDS, terminators) were cloned using Gibson assembly into starting plasmids (Level 0), with flanking Bsa1 cut sites. DNA parts with repeated sequences, such as some of the synthetic promoters, were generated by stitching together oligonucleotides using type IIS restriction enzymes and T4-DNA ligases. Cassettes composed of promoters, CDS, and terminators were assembled using Golden Gate reactions to generate constructs (Level 1) in which entire expression cassettes were flanked by Bbs1 cut sites. Various level 1 cassettes were then combined using Bbs1-based Golden Gate cloning into the binary vector pJAB2155. The sequences for all plasmids used in this study are provided as Supplementary Data.

### Transient expression assays

*N. benthamiana* plants were infiltrated with *A. tumefaciens* as described previously with a few modifications to enable fast and reliable quantitative measurements of gene expression. Briefly, *A. tumefaciens* cultures were inoculated from frozen into 14-ml culture tubes (Fisher Scientific 352059) containing 2 mL of LB media with antibiotics. Cultures were grown for 24 h at 28°C and 250 r.p.m. The next day, cultures were back diluted to an OD600 of 0.02 in fresh LB media + antibiotics and grown for another 24 h at 28°C and 250 r.p.m. Next, cells were collected by centrifugation (8 min, 3,220 rpm), resuspended in 2 mL infiltration buffer (10 mM MgCl_2_ (Sigma M4880), 10 mM MES pH 5.7 (Sigma M8250), 150 μM acetosyringone (Sigma Aldrich D134406)), and incubated at room temperature for ~3 h. Agrobacterium strains were then mixed to generate samples for leaf infiltration. In each sample, strains containing circuit components were added to the mixture at an OD600 of 0.3 and an Agrobacterium strain containing the P19-silencing suppressor was added at an OD600 of 0.2. Final sample volumes were adjusted to 1 mL using infiltration buffer. Samples were infiltrated into the abaxial side of the *N. benthamiana* leaves using a 1 mL needleless syringe (Fisher Scientific 22-253-260). Infiltrated plants were returned to normal growth conditions for ~72h.

To enable reliable quantitative measurements of synthetic parts and reduce the impact of expression variation within and between leaves, all GFP measurements were normalized to a constitutively expressed mCherry encoded on the same plasmid. mCherry was expressed using the Arabidopsis *UBIQUITIN 10* promoter (*proUBQ10::mCherry*). Throughout the paper, we report the ratio of GFP to mCherry. Raw GFP and mCherry fluorescence data is provided as Supplementary Data.

To prevent Agrobacterium from expressing *GFP* and distorting fluorescence measurements, we introduced an intron to the *GFP* gene (Supplementary Fig. 1). In initial experiments and previous studies, Agrobacterium was capable of using some plant promoters to express *GFP*. Addition of the potato ST-LS1 intron to GFP eliminated Agrobacterium expression, while leaving plant expression unaffected (Supplementary Fig. 1).

When measuring activity of the synthetic TFs and promoters in *Nicotiana benthamiana* leaves, Agrobacterium strains containing plasmids that encoded a synthetic promoter are infiltrated alone or mixed with strains containing plasmids that encoded a synthetic transcription factor under the control of the constitutively active *G0190* promoter (41). GFP fluorescence was normalized by *mCherry* encoded on the same plasmid as *GFP*. For orthogonality measurements, the synthetic promoter and transcription factor expression cassettes were combined into single plasmids.

*N. benthamiana* leaf images were collected using a fluorescence stereo microscope (Leica Microsystems THUNDER Imager Model Organism). GFP and mCherry images were collected using the following filter sets: GFP excitation 450 - 490 nm, GFP emission 500 – 550 nm, mCherry excitation 530 – 560 nm, mCherry emission 590 - 650 nm. The gain and exposure times were held constant at 2.0 and 600 ms, respectively. Leica LasX Navigator software was used to stitch individual frames together to generate whole-leaf images.

Quantitative measurements of *N. benthamiana* fluorescence were obtained using a Spark Multimode Microplate Reader (Tecan). Four leaf disks (5.5 mm diameter) were harvested per leaf using the back of a pipette tip (P20 Rainin 17005873) and placed into the bottom 96 well plates (Thomas Scientific 655096) using forceps. Each well contained 5 μL of water to adhere leaf discs to the bottom of the well. Leaf disks were placed into the plates abaxial side down. GFP and mCherry measurements were performed using the following settings: GFP excitation 408 nm, GFP emission 508 nm, GFP bandwidth 5 nm, mCherry excitation 587 nm, mCherry emission 610 nm, mCherry bandwidth 5 nm. Both fluorescence measurements were made from the bottom to capture fluorescence from the abaxial site of the leaf.

### Confocal microscopy for circuit characterization in Arabidopsis

For defining spatial patterns of expression in root tips, *A. thaliana* seeds were germinated and grown on standard MS medium. Five days after sowing, roots were stained with 10 μg/mL propidium iodide (PI – to visualize cell outlines) (Invitrogen P1304MP) for 5 min, mounted on slides, and imaged on a Leica SP8 confocal microscope. Excitation of GFP and PI was performed using lasers at 488 nm excitation with 500 – 550 nm emission filter and 488 nm excitation with 597 –664 nm emission filter, respectively. The gain and 488 nm laser power were held constant at 100 and 30% for every root unless otherwise indicated. Root tips were imaged at the midline to visualize the quiescent center and all surrounding root cell types. We screened eight T1 seedlings per genotype for expression patterns that match the expected gate output. Four plants with the highest expression were transplanted to soil after imaging and allowed to set seed. Three lines with a single-copy T-DNA insertion were imaged in T2 to generate the images reported in this manuscript.

### Root phenotyping

For analysis of root length and root branch density, red fluorescent T1 seeds were germinated and grown on standard MS media. Ten days after sowing, the plates were imaged using a dual light flatbed scanner (Epson V800). Total root length was quantified using the ImageJ segmented line tool and measured from the hypocotyl-root junction to the root tip. The number of emerged lateral roots were counted from the scanned images. Root density calculated as the number of emerged lateral roots divided by total primary root length. Root hair images are a 0.5 cm segment of the primary root, one cm above root tip.

For analysis of root gravitropism, T2 seeds were germinated and grown on standard MS media. Five days after sowing, root structure was imaged by scanning whole plates using a dual light flatbed scanner (Epson V800). *slr-1-GFP* expression was imaged using a confocal microscope as described above. Root tip organization was imaged using confocal brightfield. Four independent lines with strong agravitropic phenotypes in the T1 generation were selected for propagation to T2 and subsequent phenotypic analysis.

## Acknowledgments

We thank Tom Beekman for the Arabidopsis *slr* mutant and Tamara Vellosillo for assistance in establishing imaging protocols.

## Funding

Burroughs Wellcome Fund (JANB)

Chan Zuckerburg Biohub (JANB)

U.S. Department of Energy Biological and Environmental Research program grant DE-

SC0008769 (JRD)

Faculty Scholar grant from Howard Hughes Medical Institute and the Simons Foundation 55108515 (JRD)

## Author contributions

Conceptualization: JANB, JRD

Data acquisition: JANB, KJM, KK

Visualization: JANB

Funding acquisition: JANB, JRD

Project administration: JRD

Supervision: JANB, JRD

Writing – original draft: JANB, JRD

Writing – review & editing: JANB, JRD

## Competing interests

Authors declare that they have no competing interests.

## Data and materials availability

All plasmid materials and bacterial strains will be madde available through AddGene. Sequence files and raw data are available as supplementary materials.

## Notes

### Competing Interest Statement

The authors have declared no competing interest.

## References

1. S. J. Gurr, P. J. Rushton, Engineering plants with increased disease resistance: what are we going to express? Trends in Biotechnology. 23, 275–282 (2005).

2. B. Wang, J. Li, Understanding the Molecular Bases of Agronomic Trait Improvement in Rice[OPEN]. Plant Cell. 31, 1416–1417 (2019).

3. R. Shaw, X. Tian, J. Xu, Single-Cell Transcriptome Analysis in Plants: Advances and Challenges. Molecular Plant. 14, 115–126 (2021).

4. A. E. Richardson, J. Cheng, R. Johnston, R. Kennaway, B. R. Conlon, A. B. Rebocho, H. Kong, M. J. Scanlon, S. Hake, E. Coen, Evolution of the grass leaf by primordium extension and petiole-lamina remodeling. Science (2021), doi:10.1126/science.abf9407.

5. S. Ali, W.-C. Kim, A Fruitful Decade Using Synthetic Promoters in the Improvement of Transgenic Plants. Frontiers in Plant Science. 10, 1433 (2019).

6. L. Laplaze, B. Parizot, A. Baker, L. Ricaud, A. Martinière, F. Auguy, C. Franche, L. Nussaume, D. Bogusz, J. Haseloff, GAL4-GFP enhancer trap lines for genetic manipulation of lateral root development in Arabidopsis thaliana. J Exp Bot. 56, 2433–2442 (2005).

7. J. Bai, X. Wang, H. Wu, F. Ling, Y. Zhao, Y. Lin, R. Wang, Comprehensive construction strategy of bidirectional green tissue-specific synthetic promoters. Plant Biotechnology Journal. 18, 668–678 (2020).

8. R. Wang, M. Zhu, R. Ye, Z. Liu, F. Zhou, H. Chen, Y. Lin, Novel green tissue-specific synthetic promoters and cis-regulatory elements in rice. Sci Rep. 5, 18256 (2015).

9. L. Chen, Y. Li, Y. Wang, W. Li, X. Feng, L. Zhao, Use of High Resolution Spatiotemporal Gene Expression Data to Uncover Novel Tissue-Specific Promoters in Tomato. Agriculture. 11, 1195 (2021).

10. A. A. K. Nielsen, B. S. Der, J. Shin, P. Vaidyanathan, V. Paralanov, E. A. Strychalski, D. Ross, D. Densmore, C. A. Voigt, Genetic circuit design automation. Science. 352 (2016), doi:10.1126/science.aac7341.

11. M. Taketani, J. Zhang, S. Zhang, A. J. Triassi, Y.-J. Huang, L. G. Griffith, C. A. Voigt, Genetic circuit design automation for the gut resident species Bacteroides thetaiotaomicron. Nat Biotechnol. 38, 962–969 (2020).

12. D. Mishra, T. Bepler, B. Teague, B. Berger, J. Broach, R. Weiss, An engineered protein-phosphorylation toggle network with implications for endogenous network discovery. Science (2021), doi:10.1126/science.aav0780.

13. X. J. Gao, L. S. Chong, M. S. Kim, M. B. Elowitz, Programmable protein circuits in living cells. Science (2018), doi:10.1126/science.aat5062.

14. K. A. Schaumberg, M. S. Antunes, T. K. Kassaw, W. Xu, C. S. Zalewski, J. I. Medford, A. Prasad, Quantitative characterization of genetic parts and circuits for plant synthetic biology. Nat Methods. 13, 94–100 (2016).

15. T. K. Kassaw, A. J. Donayre-Torres, M. S. Antunes, K. J. Morey, J. I. Medford, Engineering synthetic regulatory circuits in plants. Plant Science. 273, 13–22 (2018).

16. A. J. Meyer, T. H. Segall-Shapiro, E. Glassey, J. Zhang, C. A. Voigt, Escherichia coli “Marionette” strains with 12 highly optimized small-molecule sensors. Nat Chem Biol. 15, 196–204 (2019).

17. B. C. Stanton, A. A. K. Nielsen, A. Tamsir, K. Clancy, T. Peterson, C. A. Voigt, Genomic mining of prokaryotic repressors for orthogonal logic gates. Nat Chem Biol. 10, 99–105 (2014).

18. I. Sadowski, J. Ma, S. Triezenberg, M. Ptashne, GAL4-VP16 is an unusually potent transcriptional activator. Nature. 335, 563–564 (1988).

19. C. Dingwall, R. A. Laskey, Nuclear targeting sequences — a consensus? Trends in Biochemical Sciences. 16, 478–481 (1991).

20. R. X. Fang, F. Nagy, S. Sivasubramaniam, N. H. Chua, Multiple cis regulatory elements for maximal expression of the cauliflower mosaic virus 35S promoter in transgenic plants. Plant Cell. 1, 141–150 (1989).

21. P. N. Benfey, L. Ren, N. H. Chua, Tissue-specific expression from CaMV 35S enhancer subdomains in early stages of plant development. EMBO J. 9, 1677–1684 (1990).

22. J. T. Odell, F. Nagy, N.-H. Chua, Identification of DNA sequences required for activity of the cauliflower mosaic virus 35S promoter. Nature. 313, 810–812 (1985).

23. J. B. Lucks, L. Qi, W. R. Whitaker, A. P. Arkin, Toward scalable parts families for predictable design of biological circuits. Current Opinion in Microbiology. 11, 567–573 (2008).

24. J. Li, R. Blue, B. Zeitler, T. L. Strange, J. R. Pearl, D. H. Huizinga, S. Evans, P. D. Gregory, F. D. Urnov, J. F. Petolino, Activation domains for controlling plant gene expression using designed transcription factors. Plant Biotechnology Journal. 11, 671–680 (2013).

25. E. Pierre-Jerome, S. S. Jang, K. A. Havens, J. L. Nemhauser, E. Klavins, Recapitulation of the forward nuclear auxin response pathway in yeast. PNAS. 111, 9407–9412 (2014).

26. M. Ohta, K. Matsui, K. Hiratsu, H. Shinshi, M. Ohme-Takagi, Repression Domains of Class II ERF Transcriptional Repressors Share an Essential Motif for Active Repression. The Plant Cell. 13, 1959–1968 (2001).

27. A. Pérez-González, E. Caro, Benefits of using genomic insulators flanking transgenes to increase expression and avoid positional effects. Sci Rep. 9, 8474 (2019).

28. M. Kamiya, S.-Y. Higashio, A. Isomoto, J.-M. Kim, M. Seki, S. Miyashima, K. Nakajima, Control of root cap maturation and cell detachment by BEARSKIN transcription factors in Arabidopsis. Development. 143, 4063–4072 (2016).

29. M. M. Marquès-Bueno, A. K. Morao, A. Cayrel, M. P. Platre, M. Barberon, E. Caillieux, V. Colot, Y. Jaillais, F. Roudier, G. Vert, A versatile Multisite Gateway-compatible promoter and transgenic line collection for cell type-specific functional genomics in Arabidopsis. The Plant Journal. 85, 320–333 (2016).

30. I. Blilou, J. Xu, M. Wildwater, V. Willemsen, I. Paponov, J. Friml, R. Heidstra, M. Aida, K. Palme, B. Scheres, The PIN auxin efflux facilitator network controls growth and patterning in Arabidopsis roots. Nature. 433, 39–44 (2005).

31. N. M. Clark, M. A. de Luis Balaguer, R. Sozzani, Experimental data and computational modeling link auxin gradient and development in the Arabidopsis root. Frontiers in Plant Science. 5, 328 (2014).

32. R. Rellán-Álvarez, G. Lobet, J. R. Dinneny, Environmental Control of Root System Biology. Annu Rev Plant Biol. 67, 619–642 (2016).

33. J. P. Lynch, Root phenotypes for improved nutrient capture: an underexploited opportunity for global agriculture. New Phytologist. 223, 548–564 (2019).

34. H. Fukaki, S. Tameda, H. Masuda, M. Tasaka, Lateral root formation is blocked by a gain-of-function mutation in the SOLITARY-ROOT/IAA14 gene of Arabidopsis. The Plant Journal. 29, 153–168 (2002).

35. K. Hasselt, S. Rankl, S. Worsch, A. Burkovski, Adaptation of AmtR-controlled gene expression by modulation of AmtR binding activity in Corynebacterium glutamicum. J Biotechnol. 154, 156–162 (2011).

36. R. P. Hellens, E. A. Edwards, N. R. Leyland, S. Bean, P. M. Mullineaux, pGreen: a versatile and flexible binary Ti vector for Agrobacterium-mediated plant transformation. Plant Mol Biol. 42, 819–832 (2000).

37. S. J. Clough, A. F. Bent, Floral dip: a simplified method for Agrobacterium-mediated transformation of Arabidopsis thaliana. The Plant Journal. 16, 735–743 (1998).

38. S. Emami, M. Yee, J. Dinneny, A robust family of Golden Gate Agrobacterium vectors for plant synthetic biology. Frontiers in Plant Science. 4, 339 (2013).

39. D. G. Gibson, L. Young, R.-Y. Chuang, J. C. Venter, C. A. Hutchison, H. O. Smith, Enzymatic assembly of DNA molecules up to several hundred kilobases. Nat Methods. 6, 343–345 (2009).

40. C. Engler, R. Kandzia, S. Marillonnet, A One Pot, One Step, Precision Cloning Method with High Throughput Capability. PLOS ONE. 3, e3647 (2008).

41. J. Zuo, Q.-W. Niu, N.-H. Chua, An estrogen receptor-based transactivator XVE mediates highly inducible gene expression in transgenic plants. The Plant Journal. 24, 265–273 (2000).

42. G. Vancanneyt, R. Schmidt, A. O’Connor-Sanchez, L. Willmitzer, M. Rocha-Sosa, Construction of an intron-containing marker gene: Splicing of the intron in transgenic plants and its use in monitoring early events in Agrobacterium-mediated plant transformation. Molec. Gen. Genet. 220, 245–250 (1990).

43. A. F. Ibrahim, J. A. Watters, G. P. Clark, C. J. Thomas, J. W. Brown, C. G. Simpson, Expression of intron-containing GUS constructs is reduced due to activation of a cryptic 5’ splice site. Mol Genet Genomics. 265, 455–460 (2001).

44. F. F. Assaad, E. R. Signer, Cauliflower mosaic virus P35S promoter activity in Escherichia coli. Molec. Gen. Genet. 223, 517–520 (1990).

45. D. Jacob, A. Lewin, B. Meister, B. Appel, Plant-specific promoter sequences carry elements that are recognised by the eubacterial transcription machinery. Transgenic Res. 11, 291–303 (2002).

