## Supplementary Figures for "Synthetic genetic circuits enable reprogramming of plant roots"

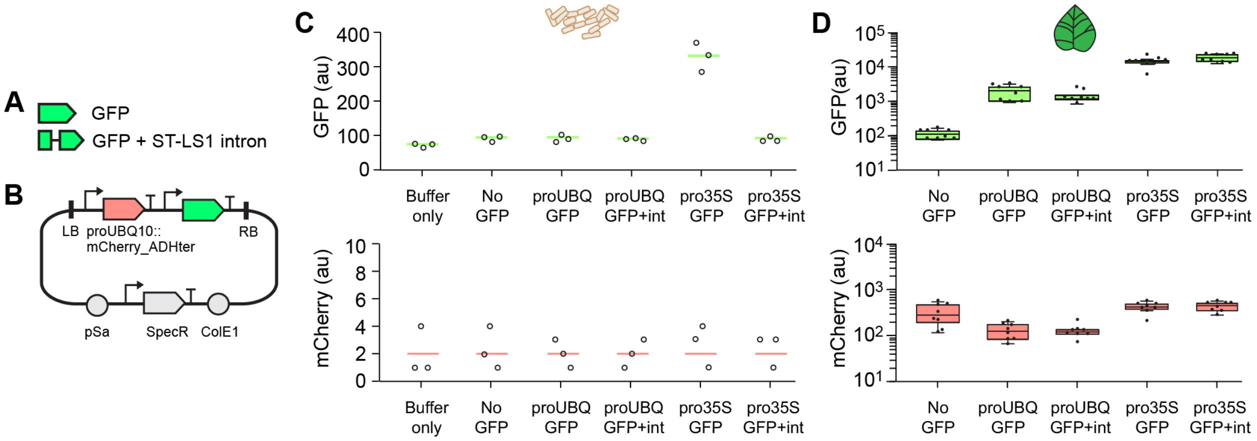


**Fig. S1. Addition of intron to GFP.** (**A**) Schematic of *GFP* and *GFP* containing intron 2 from the potato gene *ST-LS1*. This intron was previously introduced to the *uidA* gene encoding *b*-glucoronidase (GUS) to block Agrobacterium expression (42, 43). (**B**) Plasmid for measuring promoter activity. *mCherry* is constitutively expressed from the Arabidopsis *UBIQUITIN 10* promoter. *GFP* is expressed by a synthetic promoter or plant promoter. mCherry is used as an internal control to account for variation in conjugation efficiency and leaf metabolic activity (Methods). (**C**) *GFP* and *mCherry* expression in *Agrobacterium tumefaciens*. As seen in previous studies, the CaMV 35S promoter is active in Agrobacterium (42, 44, 45). Addition of an intron blocks expression of transgenes in Agrobacterium (42). The *UBIQUITIN 10* promoter is not active in Agrobacterium. Colored lines show the average of three replicates collected on different days. Individual data points shown as open circles. (**D**) *GFP* and *mCherry* expression in *Nicotiana benthamiana* leaves. Box plots show the median of twelve leaf punches collected from three leaves infiltrated and measured on different days. Box plot hinges indicate the first and third quartiles. Individual data points shown as black dots.

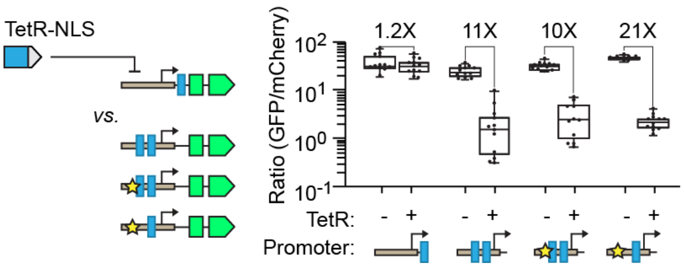


**Fig. S2. Tuning synthetic TetR-repressible promoter.** Variants of the synthetic repressible promoter that responds to TetR-NLS repressors. An eight base-pair segment of the CaMV 35S promoter (+35 to +43) was altered to increase the promoters' basal activity (ACCCTTCC changed to TAGATAAT). Modification denoted with a star. As in Figure 1, synthetic promoter activity was measured in *N. benthamiana*. TFs and promoters were encoded on separate plasmids and delivered to leaves in separate Agrobacterium strains. GFP expression was normalized by a constitutively expressed *mCherry* encoded on the same plasmid as *GFP* (Methods). All synthetic TFs were expressed using the *G0190* promoter. Box plots show the median of twelve leaf punches collected from three leaves that were infiltrated and measured on different days (four punches per leaf). Box plot hinges indicate the first and third quartiles. Letters denote significant differences in normalized GFP expression ($p < 0.01$, student's two-tailed T-test). Fold change for significantly active transcription factors was calculated by dividing the average ON state by the average OFF state.

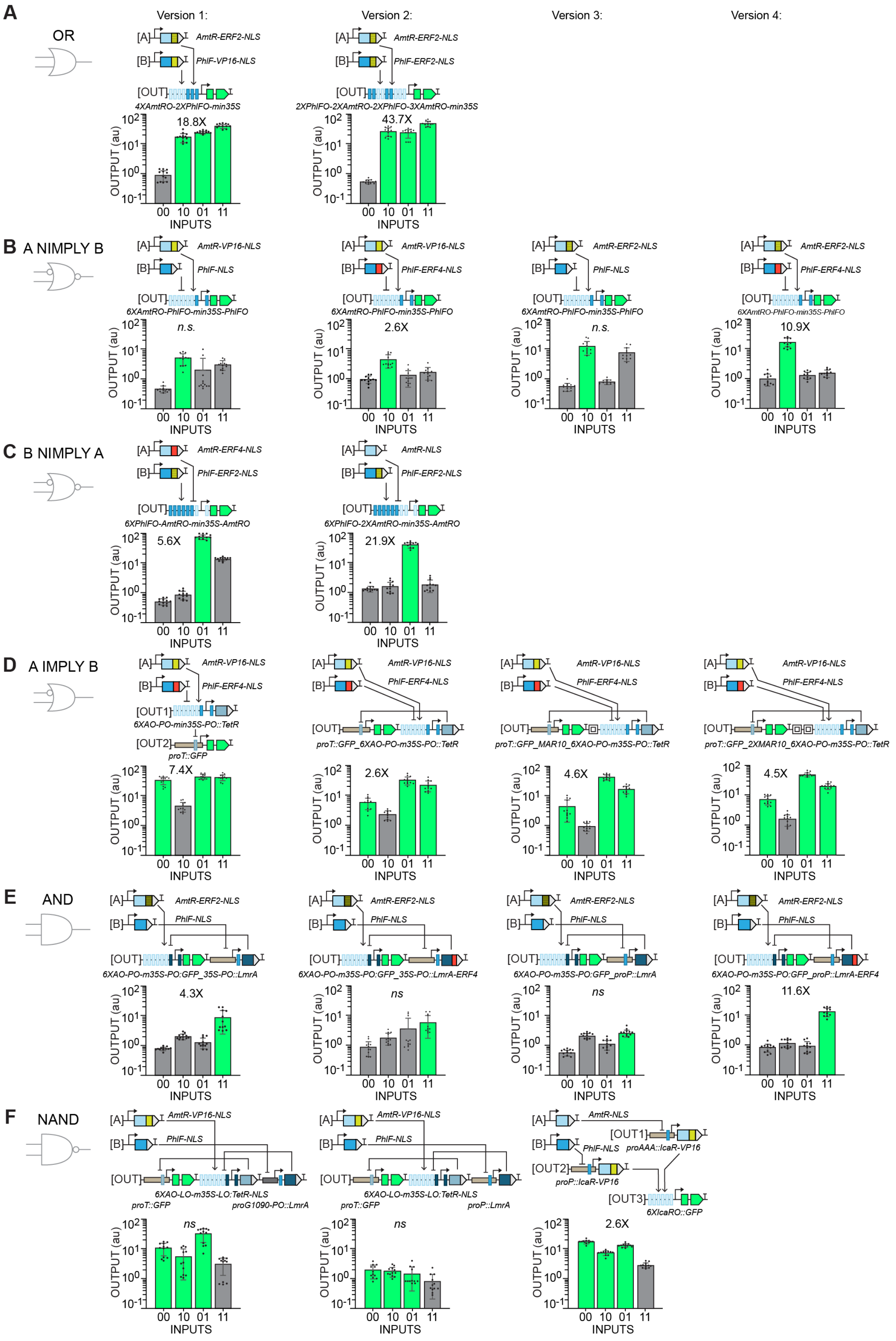
A
OR
Version 1:
Version 2:
Version 3:
Version 4:
[A] AmtR-ERF2-NLS
[B] PhlF-VP16-NLS
[OUT] 4XAmtRO-2XPhlFO-min35S
18.8X
[A] AmtR-ERF2-NLS
[B] PhlF-ERF2-NLS
[OUT] 2XPhlFO-2XAmtRO-2XPhlFO-3XAmtRO-min35S
43.7X
B A NIMPLY B
[A] AmtR-VP16-NLS
[B] PhlF-NLS
[OUT] 6XAmtRO-PhlFO-min35S-PhlFO
n.s.
[A] AmtR-VP16-NLS
[B] PhlF-ERF4-NLS
[OUT] 6XAmtRO-PhlFO-min35S-PhlFO
2.6X
[A] AmtR-ERF2-NLS
[B] PhlF-NLS
[OUT] 6XAmtRO-PhlFO-min35S-PhlFO
n.s.
[A] AmtR-ERF2-NLS
[B] PhlF-ERF4-NLS
[OUT] 6XAmtRO-PhlFO-min35S-PhlFO
10.9X
C B NIMPLY A
[A] AmtR-ERF4-NLS
[B] PhlF-ERF2-NLS
[OUT] 6XPhlFO-AmtRO-min35S-AmtRO
5.6X
[A] AmtR-NLS
[B] PhlF-ERF2-NLS
[OUT] 6XPhlFO-2XAmtRO-min35S-AmtRO
21.9X
D A IMPLY B
[A] AmtR-VP16-NLS
[B] PhlF-ERF4-NLS
[OUT1] 6XAO-PO-min35S-PO::TetR
[OUT2] proT::GFP
7.4X
[A] AmtR-VP16-NLS
[B] PhlF-ERF4-NLS
[OUT] proT::GFP_6XAO-PO-m35S-PO::TetR
2.6X
[A] AmtR-VP16-NLS
[B] PhlF-ERF4-NLS
[OUT] proT::GFP_MAR10_6XAO-PO-m35S-PO::TetR
4.6X
[A] AmtR-VP16-NLS
[B] PhlF-ERF4-NLS
[OUT] proT::GFP_2XMAR10_6XAO-PO-m35S-PO::TetR
4.5X
E AND
[A] AmtR-ERF2-NLS
[B] PhlF-NLS
[OUT] 6XAO-PO-m35S-PO:GFP_35S-PO::LmrA
4.3X
[A] AmtR-ERF2-NLS
[B] PhlF-NLS
[OUT] 6XAO-PO-m35S-PO:GFP_35S-PO::LmrA-ERF4
ns
[A] AmtR-ERF2-NLS
[B] PhlF-NLS
[OUT] 6XAO-PO-m35S-PO:GFP_proP::LmrA
ns
[A] AmtR-ERF2-NLS
[B] PhlF-NLS
[OUT] 6XAO-PO-m35S-PO:GFP_proP::LmrA-ERF4
11.6X
F NAND
[A] AmtR-VP16-NLS
[B] PhlF-NLS
[OUT] 6XAO-LO-m35S-LO:TetR-NLS
proT::GFP
proG1090-PO::LmrA
ns
[A] AmtR-VP16-NLS
[B] PhlF-NLS
[OUT] 6XAO-LO-m35S-LO:TetR-NLS
proT::GFP
proP::LmrA
ns
[A] AmtR-NLS
[B] PhlF-NLS
[OUT1] proAAA::IcaR-VP16
[OUT2] proP::IcaR-VP16
[OUT3] 6XIcaRO::GFP
2.6X
OUTPUT (au)
INPUTS
00 10 01 11

**Fig. S3. Tuning logic gate behavior in *N. benthamiana*.** (A) OR gate variants. The number and location of AmtR and PhlF operators were varied in the output promoter. (B) A NIMPLY B gate variants. Different combinations of AmtR and PhlF-based synthetic TFs were tested. (C) A NIMPLY B gate variants. Different combinations of AmtR and PhlF-based synthetic TFs were tested and the number of AmtR operators in the output promoter was increased. (D) A IMPLY B gate variants. Arabidopsis *MATRIX ATTACHMENT REGION 10 (AtMAR10)* was tested as an insulator between expression cassettes on the output promoter. (E) AND gate variants. The promoter and synthetic TF in this gate's intermediate layer were varied. (F) NAND gate variants. In the first two versions, the PhlF-responsive promoter in this gate's intermediate layer were varied. In the third version, NAND logic was implemented using multiple repressible promoters. In all panels, circuit activity was measured in *N. benthamiana*. TFs and promoters were encoded on separate plasmids and delivered to leaves in separate Agrobacterium strains. GFP expression was normalized by a constitutively expressed *mCherry* encoded on the same plasmid as *GFP* (Methods). All synthetic TFs were expressed using the *G0190* promoter. Bar charts show the average of twelve leaf punches collected from three leaves that were infiltrated and measured on different days (four punches per gate state leaf). Error bars show standard deviation. Fold change for each gate that achieves a significant difference between ON and OFF was calculated by dividing the lowest ON state by the highest OFF state ($p < 0.01$, student's two-tailed T-test). *N.s.* is written above gates in which the lowest ON and highest OFF states are not significantly different. Schematic colors are the same as in Fig. 1.

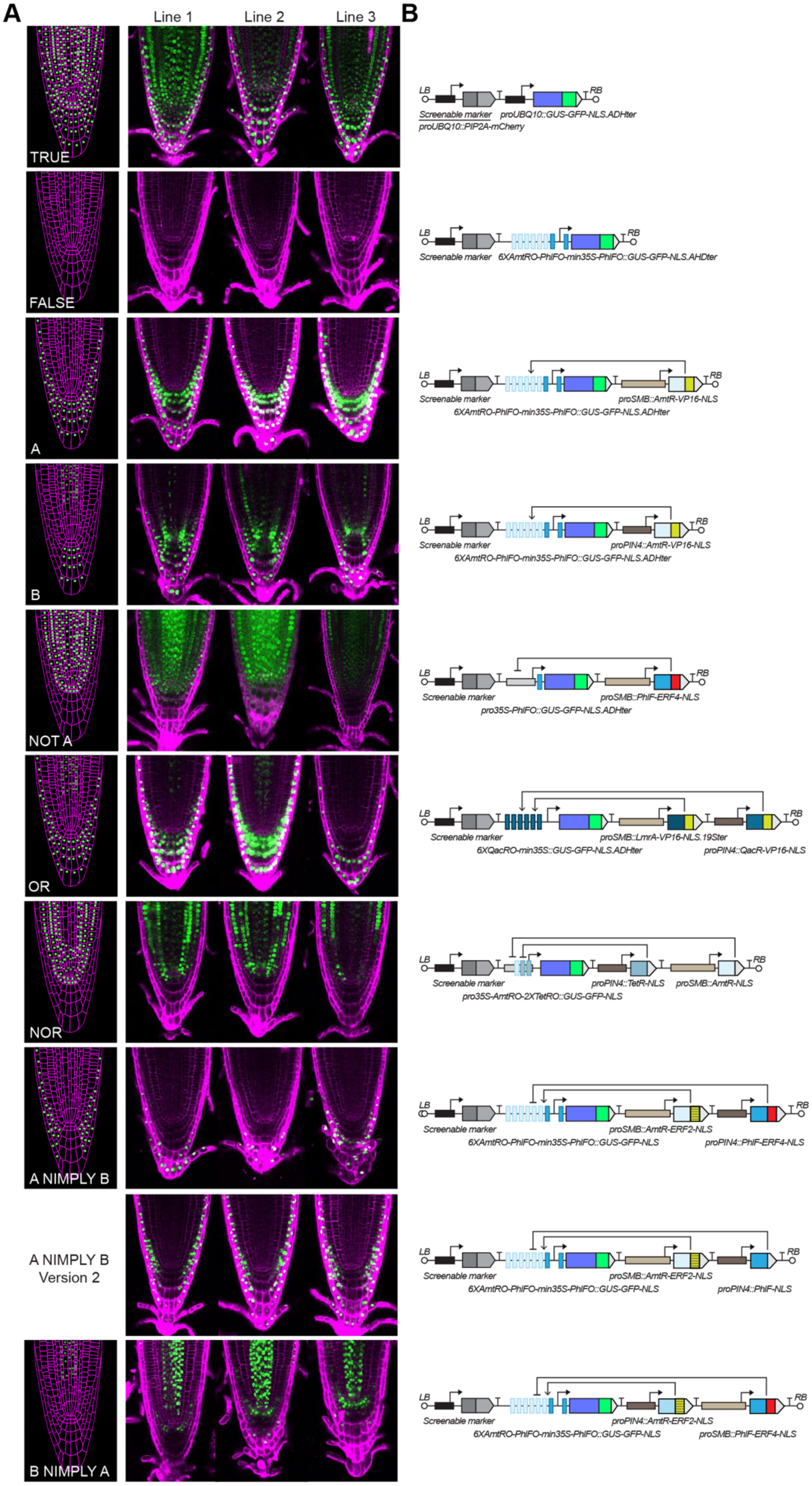

A
Line 1
Line 2
Line 3
B
TRUE
FALSE
A
B
NOT A
OR
NOR
A NIMPLY B
A NIMPLY B Version 2
B NIMPLY A
LB
RB
Screenable marker
proUBQ10::PIP2A-mCherry
proUBQ10::GUS-GFP-NLS.ADHter
6XAmtRO-PhlFO-min35S-PhlFO::GUS-GFP-NLS.AHDter
6XAmtRO-PhlFO-min35S-PhlFO::GUS-GFP-NLS.ADHter
proSMB::AmtR-VP16-NLS
proPIN4::AmtR-VP16-NLS
pro35S-PhlFO::GUS-GFP-NLS.ADHter
proSMB::PhlF-ERF4-NLS
6XQacRO-min35S::GUS-GFP-NLS.ADHter
proSMB::LmrA-VP16-NLS.19Ster
proPIN4::QacR-VP16-NLS
pro35S-AmtRO-2XTetRO::GUS-GFP-NLS
proPIN4::TetR-NLS
proSMB::AmtR-NLS
6XAmtRO-PhlFO-min35S-PhlFO::GUS-GFP-NLS
proSMB::AmtR-ERF2-NLS
proPIN4::PhlF-ERF4-NLS
proPIN4::PhlF-NLS
proPIN4::AmtR-ERF2-NLS
proSMB::PhlF-ERF4-NLS

**Fig. S4. Additional independent lines and schematics of logic gate expression patterns in Arabidopsis roots. (A)** Expected expression patterns (left) and confocal images (right) of each logic gate in the Arabidopsis root tip. Confocal images showing the expression pattern created in three independent lines. All images are of segregating T2 lines taken five days after sowing. Laser power and image processing was identical for every image to show differences in expression intensity across circuits and independent lines. **(B)** Schematics of the T-DNAs transformed into Arabidopsis for each circuit. Left (LB) and right border (RB) denote the ends of the DNA transformed into Arabidopsis. Constitutively expressed mCherry (*proUBQ10::PIP2A-mCherry*) was the screenable marker used to identify transformed seeds. Sequences of the T-DNAs available as supplementary material.

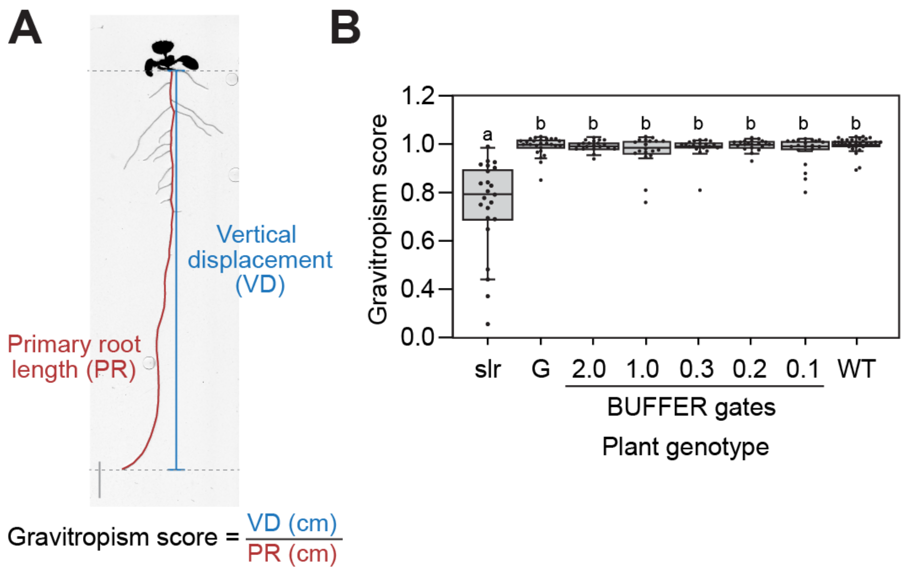


**Fig. S5. Root gravitropism in engineered plants.** (**A**) Schematic showing how root gravitropism is measured. Image is of wild type Arabidopsis, Columbia ecotype (Col-0). (**B**) Graivtropism scores of plants from Figure 4. Samples are labeled as follows: *slr-1* (slr), *proGATA23::slr* (G), wild type (WT). Box plots show the median of at least 20 T1 plants. Box plot hinges indicate the first and third quartiles. Letters denote significant differences in gravitropsim ($p < 0.01$, Student's two-tailed t-test). All measurements and images were taken ten days after sowing.

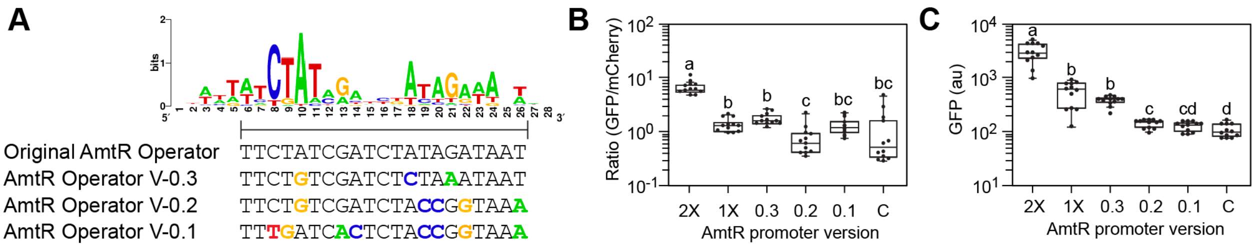


**Fig. S6. AmtR Operator (AmtRO) variants.** (**A**) The AmtR operator sequence and engineered variants. An AmtR motif logo was calculated by Hasselt *et al*. using AmtR binding sites in the *Corenybacterium glutamicum* genome (37). The height of letters represents the frequency of the corresponding nucleotides in AmtR binding sites. Black bar shows the region of the binding site used in this study as the AmtR operator. This consensus sequence was selected by a previous study that used AmtR to build synthetic genetic circuits in *Escherichia coli* (19). Consensus operator sequence and variants created for this study listed below the sequence logo. Mutations relative to the original operator are colored and bolded. (**B**) Activity of synthetic promoters containing either one or two of the original AmtR operators (promoters 1 and 2, respectively) or one of the mutant AmtR operators described in (a) (V-0.1, V-0.2, V-0.3). Promoters driving expression of GFP were co-delivered to *N. benthamiana* leaves with AmtR-ERF2 activators. Synthetic promoter activity was reported as the ratio of GFP to a constitutively expressed mCherry encoded on the same plasmid. A plasmid with no GFP gene was measured as a control (C). (**C**)_Un-normalized GFP expression by the AmtR activatable promoter variants reported in (b). In all panels, AmtR-ERF2 TFs were expressed using the *G0190* promoter (36). Box plots show the median of twelve leaf punches collected from three leaves that were infiltrated and measured on different days (four punches per leaf). Box plot hinges indicate the first and third quartiles. Black dots represent individual data points. Letters denote significant differences in normalized GFP expression ($p < 0.01$, b) or un-normalized GFP expression ($p < 0.01$, c)

(student's two-tailed T-test). Raw GFP and mCherry measurements provided as Supplementary Data.

**Supplementary Data S1.** Raw GFP and mCherry measurements for all *Nicotiana benthamiana* leaf assays.
